# Infectious virus in exhaled breath of symptomatic seasonal influenza cases from a college community

**DOI:** 10.1101/194985

**Authors:** Jing Yan, Michael Grantham, Jovan Pantelic, P. Jacob Bueno de Mesquita, Barbara Albert, Fengjie Liu, Sheryl Ehrman, Donald K. Milton, for the EMIT Consortium

**Author notes:** Corresponding author: Donald K Milton, MD, DrPH, MIAEH Rm 2234V, School of Public Health, 4200 Valley Dr., College Park, MD, 20742. Membership of the EMIT Consortium is provided in the Supporting Information.

## Abstract

Little is known about the amount and infectiousness of influenza virus shed into exhaled breath. This contributes to uncertainty about the importance of airborne influenza transmission. We screened 355 symptomatic volunteers with acute respiratory illness and report 142 cases with confirmed influenza infection who provided 218 paired nasopharyngeal (NP) and 30-minute breath samples (coarse >5 μm and fine ≤5 μm fractions) on days 1 to 3 post symptom onset. We assessed viral RNA copy number for all samples and cultured NP swabs and fine aerosols. We recovered infectious virus from 52 (39%) of the fine aerosols and 150 (89%) of the NP swabs with valid cultures. The geometric mean RNA copy numbers were 3.8×10^4^/30-min fine, 1.2×10^4^/30-min coarse aerosol sample, and 8.2×10^8^ per NP swab. Fine and coarse aerosol viral RNA was positively associated with body mass index (fine p<0.05, coarse p<0.10) and number of coughs (fine p<0.001, coarse p<0.01) and negatively associated with increasing days since symptom onset (fine p<0.05 to p<0.01, coarse p<0.10) in adjusted models. Fine aerosol viral RNA was also positively associated with having influenza vaccination for both the current and prior season (p<0.01). NP swab viral RNA was positively associated with upper respiratory symptoms (p<0.01) and negatively associated with age (p<0.01) but was not significantly associated with fine or coarse aerosol viral RNA or their predictors. Sneezing was rare, and sneezing and coughing were not necessary for infectious aerosol generation. Our observations suggest that influenza infection in the upper and lower airways are compartmentalized and independent.

**Significance:** Lack of human data on influenza virus aerosol shedding fuels debate over the importance of airborne transmission. We provide overwhelming evidence that humans generate infectious aerosols and quantitative data to improve mathematical models of transmission and public health interventions. We show that sneezing is rare and not important for, and that coughing is not required for influenza virus aerosolization. Our findings, that upper and lower airway infection are independent and that fine particle exhaled aerosols reflect infection in the lung, open a new pathway for understanding the human biology of influenza infection and transmission. Our observation of an association between repeated vaccination and increased viral aerosol generation demonstrated the power of our method, but needs confirmation.

## Introduction

The nature of infectious contacts and the relative importance of contact, large droplet spray and aerosol (droplet nuclei) transmission remain controversial.(1–6) Non-pharmaceutical interventions have been employed to control and reduce the impact of influenza epidemics and pandemics.(7) However, to design effective non-pharmaceutical interventions, it is necessary to accurately define the relative and absolute contribution of each route of transmission(8) and implement interventions that impede those of principal importance.

Mathematical models that have been used to understand and estimate the contribution of each mode are very sensitive to estimates of unmeasured parameters,(9, 10) such as the viral load in exhaled breath and coughs and the frequency of sneezing by influenza cases.(8) However, due to limitations inherent to sampling virus shedding via various routes from infected individuals and the difficulty of distinguishing routes of transmission in observational studies, the quantitative dynamics and relative contributions of each route remain elusive.(4, 8) Recent reports have shown that infectious influenza virus can be recovered from exhaled aerosols.(11–13) These studies, based on small numbers of cases or artificial breathing maneuvers, do not provide sufficient data to quantify the extent of aerosol shedding during natural breathing, identify the contributions of spontaneous coughs and sneezes commonly thought to be the most important mechanism for viral shedding, nor identify other factors that may impact viral aerosol shedding. We address these key knowledge gaps by characterizing influenza virus in exhaled breath from community acquired influenza cases during natural breathing, prompted speech, coughing, and sneezing, and assess the infectivity of naturally occurring influenza aerosols.

## Results

We screened 355 volunteers with acute respiratory illness; the 178 volunteers who met enrollment criteria provided 278 visits for sample collection. We confirmed influenza infection in 156 (88%) of the enrolled participants using RT-qPCR; 152 had at least one positive NP swab and 4 (3%) were confirmed based on positive aerosol samples alone. NP swab analysis was positive for 8 (33%) of 24 randomly selected volunteers from among the 177 screened who did not meet enrollment criteria; thus, sensitivity and specificity of our enrollment criteria, during the 2012–13 season, were approximately 73% (95% confidence interval (CI) 62 to 84%) and 84% (95% CI 80 to 88%), respectively. In the reported analyses, we excluded eight visits made on the day of symptom onset, 10 made >3 days after onset, seven with missing data for cough, and three with incomplete RT-qPCR data (Supplementary Table S1 and Fig. S1). The resulting data set for confirmed cases with complete data on RNA copies, cough, and symptoms included 218 visits by 142 cases: 89 influenza A (83 H3, 3 pdmH1, 3 unsubtypable), 50 influenza B, and 3 dual influenza infection cases.

Our study population (Table 1) consisted mostly of young adults (19–21 years) with a high asthma prevalence (21%), normal Body Mass Index (BMI, median=22.7; 7% underweight, 20% overweight, and 8% obese, Table S2), and a low self-reported influenza vaccination rate (22%). We observed at least one cough during 195 (89%) and one or more sneezes during 11 (5%) of the 218 visits. Cough frequency varied considerably from 5 per 30 min. at the 25^th^ centile to 39 per 30 min at the 75^th^. Most volunteers rated their upper respiratory symptoms as mild to moderate, systemic symptoms as moderate to severe, and lower respiratory symptoms as mild (Fig. 1).

**Figure 1.**
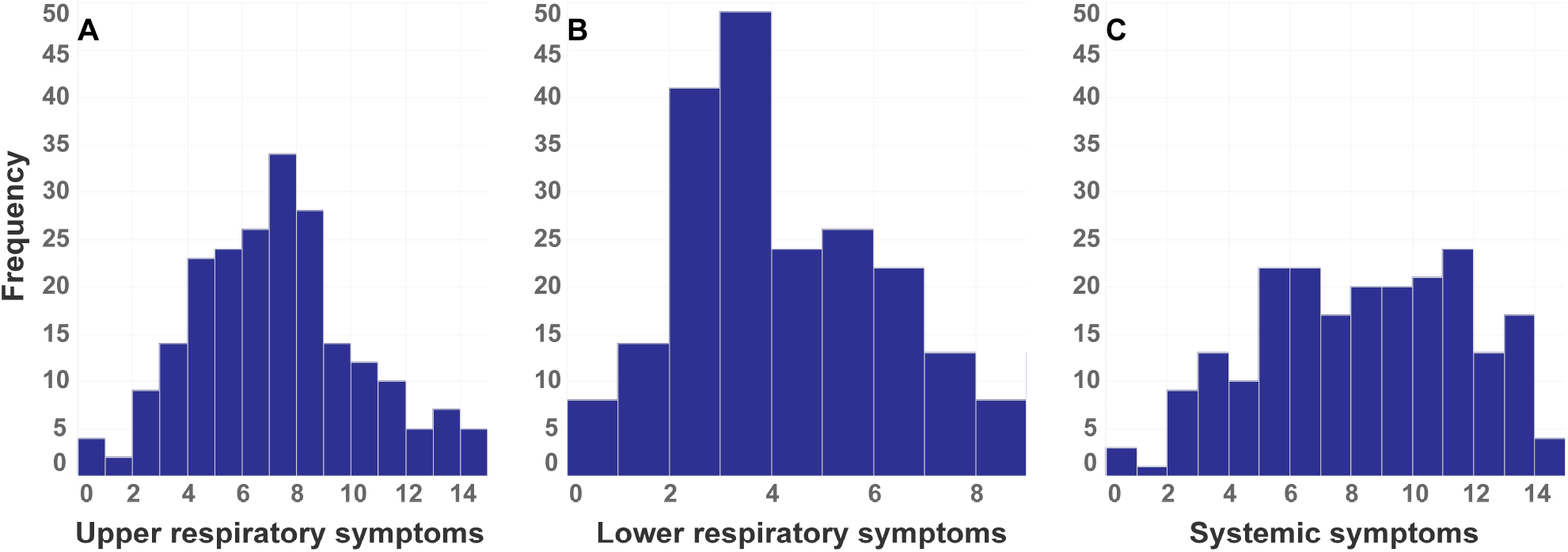
Histogram of symptom scores A) Upper respiratory symptoms (runny nose, stuffy nose, sneezing, sore throat, and earache, score range 0 to 15), B) Lower respiratory symptoms (chest tightness, shortness of breath, and cough, score range 0 to 9) and C) Systemic symptoms (malaise, headache, muscle/joint ache, fever/sweats/chills, and swollen lymph nodes, score range 0 to 15).

**Table 1.** Characteristics of Study Population

|  | Screened Only | Enrolled | Complete Data |
| --- | --- | --- | --- |
| N participants | 177 | 178 | 142 |
| Breath collection visits | 0 | 278 | 218 |
| Male (%) | 89 (50) | 91 (51) | 69 (49) |
| Influenza vaccination current season (%) | 53 (30) | 42 (24) | 31 (22) |
| Influenza vaccination previous season (%) | 77 (44) | 63 (35) | 48 (34) |
| Influenza vaccination current & previous (%) | 39 (22) | 32 (18) | 22 (15) |
| Asthma, self-reported (%) | – | 38 (21) | 30 (21) |
| Smoker, current (%) | – | 30 (17) | 21 (15) |
| Anti-viral medication last 24 hr (%) | 2 | 11 (6) | 7 (5) |
| Age (IQR) | 20 (19 - 22) | 21 (19 - 22) | 20 (19 - 21) |
| BMI (IQR) | 23.6 (21.3 - 26.2) | 23.2 (21.0 - 25.7) | 22.7 (20.9 - 25.5) |
| Body temperature measured (IQR) | 36.9 (36.8 - 37.1) | 37.2 (36.9 - 37.7) | 37.2 (36.9 - 37.6) |
| Median Coughs/30 minutes (IQR)* | – | 17 (6 - 39) | 18 (5 - 39) |
| Median Sneezes/30 minutes (IQR) | – | 0 (0-0) | 0 (0 - 0) |
| Median Upper respiratory symptoms (IQR) † | 7 (4 - 9) | 6 (5 - 8) | 7 (5 - 8) |
| Median Lower respiratory symptoms (IQR) | 2 (1 - 4) | 3 (2 - 5) | 3 (2 - 6) |
| Median Systemic symptoms (IQR) | 6 (3 - 9) | 8 (5 - 11) | 8 (5 - 11) |

IQR = interquartile range.
\* Cough, sneeze, and symptom scores are reported per visit
† Twelve symptoms were rated from 0 to 3 with maximum possible composite score of 15 for upper respiratory, 9 for lower respiratory, and 15 for systemic symptoms.

Infectious virus was recovered from 52 (39%) fine aerosol samples and 150 (89%) NP swabs (Table 2). Quantitative cultures were positive for 30% of the fine aerosol samples with a geometric mean (GM) for positive samples of 37 fluorescent focus units (FFU) per 30-min sample (Fig. 2A) and for 62% of NP swabs with GM for positive samples of 2,500. Using Tobit analysis to adjust the estimate of the GM for the presence of samples below the limit of detection we obtained GM 1.6 (95% CI 0.7 to 3.5) for fine aerosols and GM 60.6 (95% CI 22.7 to 1.6×10^2^) for NP swabs.

**Fig. 2.**
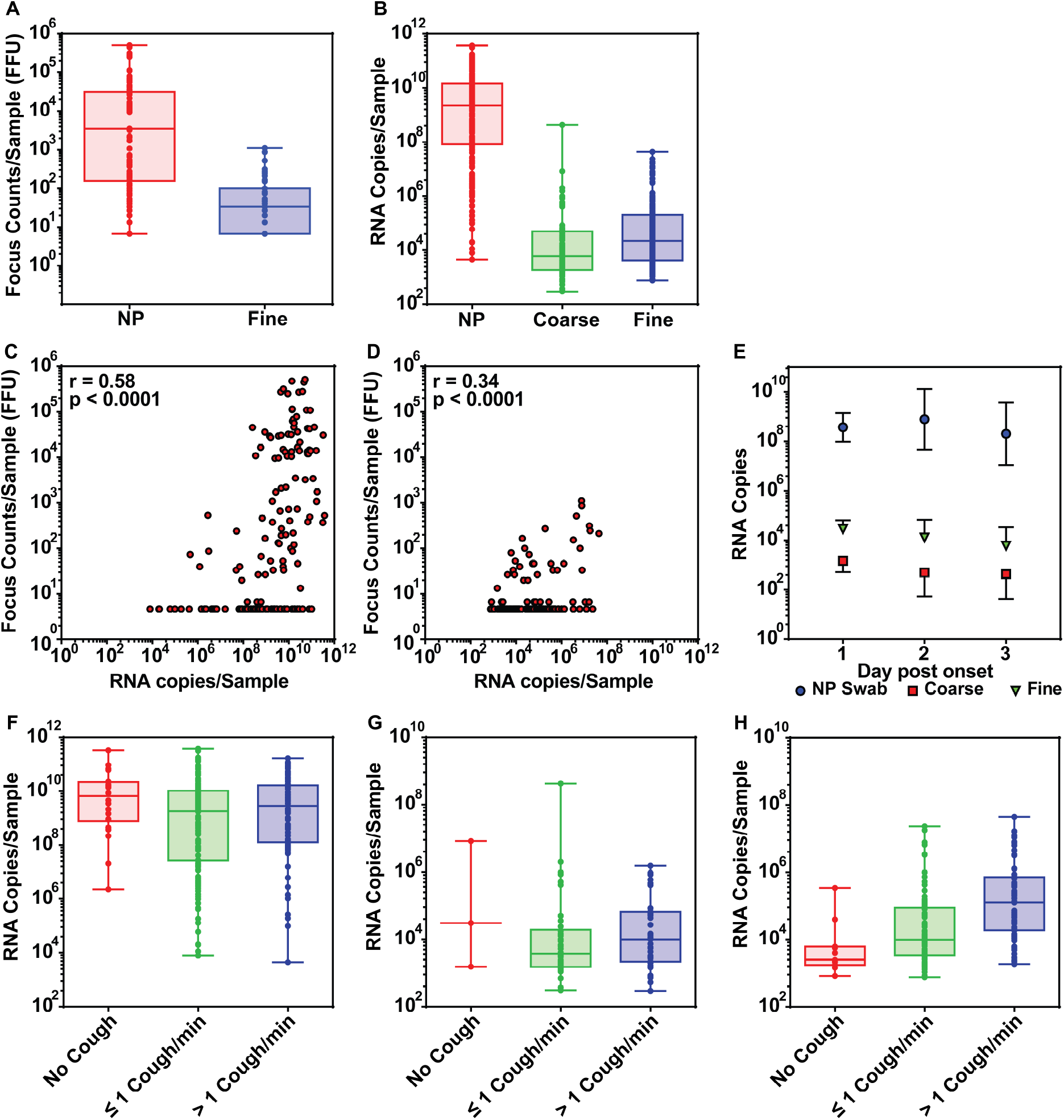
Viral shedding: (A) infectious influenza virus (fluorescent focus counts) in NP swabs and fine aerosols and (B) RNA copies in NP swabs, coarse, and fine aerosols; scatter plots and Spearman correlation coefficients of infectious virus plotted against RNA copies for (C) NP swabs and for (D) fine aerosol samples; the effect of day post symptom onset on RNA copies observed in NP swabs, coarse, and fine aerosols (E) plotted as geometric mean adjusted for missing data using Tobit analysis with error bars denoting 95% confidence intervals; and the effect of cough frequency on RNA copies observed in (F) NP swabs, (G) coarse aerosols, and (H) in fine aerosols. NP = nasopharyngeal swab, Coarse = aerosol droplets > 5 μm and Fine = aerosol droplets ≤ 5 μm in aerodynamic diameter.

**Table 2.** Viral Shedding

|  | **NP Swab** | **Coarse Aerosol** | **Fine Aerosol** |
| --- | --- | --- | --- |
| **Culture Passage** |  |  |  |
| **Valid Assays** | 169 | NA | 134 |
| **Positive (%)** | 150 (89) | NA | 52 (39) |
| **Quantitative Culture (FFU)** |  |  |  |
| **Valid Assays** | 159 | NA | 136 |
| **Positive (%)** | 98 (62) | NA | 41 (30) |
| **GM (GSD)** | $2.5 \times 10^3$ (23) | NA | 37 (4.4) |
| **Range** | ND – $5.1 \times 10^5$ | NA | ND – $1.1 \times 10^3$ |
| **RNA Copies** |  |  |  |
| **Valid Assays** | 218 | 218 | 218 |
| **Positive (%)** | 211 (97) | 88 (40) | 166 (76) |
| **GM (GSD)** | $8.2 \times 10^8$ (52) | $1.2 \times 10^4$ (14) | $3.8 \times 10^4$ (13) |
| **Range** | ND – $3.8 \times 10^{11}$ | ND – $4.3 \times 10^8$ | ND – $4.4 \times 10^7$ |

NA = Not assayed, FFU = fluorescent focus unit, GM = Geometric mean, GSD = Geometric standard deviation (only positive samples were included in computation of GM and GSD), ND = Not Detected.

Influenza virus RNA was detected in 76% of the fine aerosol samples, 40% of the coarse aerosol samples, and 97% of the NP swabs of enrolled volunteers. For the positive samples, the GM viral RNA content of fine aerosol samples was 3.8×10^4^, for coarse aerosols was 1.2×10^4^, and for NP swabs was 8.2×10^8^ (Fig. 2B). The adjusted GMs were 1.2×10^4^ (95% CI 7.0×10^3^ to 1.9×10^4^) for fine aerosols and 6.0×10^2^ (95% CI 3.0×10^2^ to 1.2×10^3^) for coarse aerosols. Quantitative culture was correlated with RNA copies in both NP swabs (Fig. 2C, r=0.58) and fine aerosols (Fig. 2D, r=0.34). The time course of shedding is shown in Fig. 2E.

Viral RNA in NP swabs was not correlated with cough frequency (number of coughs per 30 min, Fig. 2F and Fig. S2A, r=0.02). Viral RNA in in coarse aerosols was weakly correlated with cough frequency (Fig. 2G and Fig. S2B, r=0.24). However, viral RNA copy number in fine aerosols was moderately well correlated with cough frequency (Fig. 2H and Fig. S2C, r=0.45). Only 3 (13%) of 23 coarse aerosol samples where no coughs were observed had detectable viral RNA, while 11 (48%) of the corresponding 23 fine aerosol samples had detectable viral RNA and 8 were positive by culture. RNA copies in the fine aerosol, no-cough samples ranged up to 3.7×10^5^ (adjusted GM 1.5 ×10^3^, 95% CI 4.2×10^2^ to 5.3 ×10^3^) and infectious virus to 1.4×10^2^ FFU per 30-minute sample. The few sneezes observed were not associated with greater RNA copy numbers in either coarse or fine aerosols (Fig. S3).

Results of regression analyses to identify predictors of viral RNA shedding are shown in Table 3, controlled for random effects of subject and repeated observations on individuals.

**Table 3.** Predictors of Viral RNA Shedding^1^

| Parameter | NP Swab |  | Coarse Aerosol |  | Fine Aerosol |  |
| --- | --- | --- | --- | --- | --- | --- |
|  | Unadjusted | Adjusted | Unadjusted | Adjusted | Unadjusted | Adjusted |
| Age | **0.85<sup>†</sup>**<br>(**0.72 – 1.0**) | **0.78<sup>‡</sup>**<br>(**0.66 – 0.92**) | 1.0<br>(0.90 – 1.2) | 1.0<br>(0.90 – 1.1) | 0.99<br>(0.88 – 1.1) | 1.0<br>(0.90 – 1.1) |
| Male | 0.34<br>(0.14 – 2.5) | 0.68<br>(0.17 – 2.8) | 1.7<br>(0.52 – 5.6) | 1.7<br>(0.54 – 5.5) | **2.4<sup>\*</sup>**<br>(**0.88 – 6.5**) | 1.3<br>(0.39 – 4.2) |
| Asthma | 3.8<br>(0.63 – 22.9) | - | 1.38<br>(0.32 – 6.04) | - | 2.6<br>(0.77 – 8.95) | - |
| Smoker | 2.3<br>(0.30 – 18.2) | - | 0.48<br>(0.09 – 2.7) | - | 1.5<br>(0.36 – 6.1) | - |
| BMI | 1.3<br>(0.58 – 2.7) | - | **1.9<sup>\*</sup>**<br>(**0.92 – 3.98**) | 1.8<br>(0.83 – 3.8) | **1.7<sup>\*</sup>**<br>(**0.98 – 2.9**) | **2.5<sup>‡</sup>**<br>(**1.4 – 4.3**) |
| Vaccination<sup>2</sup> (current year) | 1.3<br>(0.22 – 7.6) | - | 1.4<br>(0.34 – 5.9) | - | **2.8<sup>\*</sup>**<br>(**0.85 – 9.3**) | - |
| Vaccination (previous year) | 1.55<br>(0.45 – 5.5) | - | 0.71<br>(0.15 – 3.3) | - | 2.3<br>(0.79 – 6.6) | - |
| Vaccination (current & previous) | 0.77<br>(0.10 – 5.7) | - | 1.13<br>(0.22 – 5.7) | - | **2.1<sup>‡</sup>**<br>(**1.3 – 3.2**) | **6.3<sup>‡</sup>**<br>(**1.9 – 21.5**) |
| Antiviral medication | 1.1<br>(0.03 – 33.9) | - | 0.64<br>(0.04 – 11.5) | - | 1.04<br>(0.09 – 12.0) | - |
| Influenza A | 0.67<br>(0.13 – 3.3) | - | 1.0<br>(0.18 – 5.5) | - | 0.44<br>(0.13 – 1.5) | - |
| Log (NP Swab) | - | - | 1.3<br>(0.91 – 1.82) | - | 1.2<br>(0.76 – 1.78) | - |
| Day 2 post onset | 2.13<br>(0.48 – 9.5) | - | **0.34<sup>\*</sup>**<br>(**0.10 – 1.02**) | **0.34<sup>\*</sup>**<br>(**0.10 – 1.1**) | **0.44<sup>\*</sup>**<br>(**0.18 – 1.05**) | **0.47<sup>\*</sup>**<br>(**0.20 – 1.1**) |
| Day 3 post onset | 0.55<br>(0.11 – 2.8) | - | **0.29<sup>\*</sup>**<br>(**0.08 – 1.03**) | 0.30<br>(0.08 – 1.03) | **0.20<sup>§</sup>**<br>(**0.08 – 0.52**) | **0.24<sup>‡</sup>**<br>(**0.09 – 0.59**) |
| Fever<sup>3</sup> | 3.6<br>(0.76 – 17.2) | - | 0.70<br>(0.19 – 2.7) | - | 1.7<br>(0.64 – 4.5) | - |
| Number of coughs | 1.3<br>(0.65 – 2.8) | - | **2.1<sup>‡</sup>**<br>(**1.2 – 3.8**) | **2.3<sup>‡</sup>**<br>(**1.3 – 4.0**) | **2.9<sup>¶</sup>**<br>(**1.9 – 4.6**) | **2.6<sup>§</sup>**<br>(**1.6 – 4.2**) |
| Upper respiratory symptoms | **3.4<sup>§</sup>**<br>(**1.8 – 6.6**) | **2.9<sup>‡</sup>**<br>(**1.5 – 5.7**) | 0.76<br>(0.44 – 1.3) | - | 0.88<br>(0.58 – 1.4) | - |
| Lower respiratory symptoms | **3.2<sup>\*</sup>**<br>(**0.96 – 10.8**) | 1.2<br>(0.32 – 4.8) | 0.94<br>(0.36 – 2.5) | - | 1.9<br>(0.86 – 4.1) | - |
| Systemic symptoms | **5.3<sup>‡</sup>**<br>(**1.6 – 17.7**) | 2.4<br>(0.31 – 18.3) | 0.86<br>(0.33 – 2.3) | - | 1.8<br>(0.82 – 3.7) | - |
| Male × Number of Coughs | - | - | - | - | - | **3.2<sup>†</sup>**<br>(**1.1 – 9.3**) |

1 – Effect estimates are shown as the ratio of male to female, Day 2 or Day 3 to Day 1, type A to B, yes to no vaccination, or fold increase for an interquartile range (IQR) change in age, the number of coughs, symptom reports, or BMI, or ratio of male to female coughs over the IQR (95% confidence interval for the effect estimate). All analyses are controlled for random effects of subject and sample within subject and for censoring by limit of detection using Tobit regression.
2 – Vaccination = self-reported influenza vaccination
3 – Fever = $T \geq 37.8$ measured at visit
\* $p < 0.10$ , † $p < 0.05$ , ‡ $p < 0.01$ , § $p < 0.001$ , ¶ $p < 0.0001$ from Tobit regression models with random effect of subject and sample within subject. Adjusted models were selected using the Akaike information criterion from initial models with all unadjusted parameters having $p < 0.10$ , shown in bold.

Day post symptom onset (comparing day 1 post onset with day 2 and day 3) was associated with a significant decline in viral RNA shed into fine aerosols (p < 0.05 for day 2 and p < 0.01 for day 3 in adjusted models), a borderline significant decline in coarse aerosol shedding (p < 0.10) and was not associated with a significant change in shedding detected in NP swabs (p > 0. 10).

In regression analyses, cough frequency was significantly associated with increased fine (p< 0.001 to <0.0001) and coarse (p < 0.01) aerosol shedding, but was not associated with NP shedding. Fine aerosol shedding was significantly greater for males. Analysis of an interaction of cough with sex indicated that males produced, on average, 3.2 times more virus than did females per cough. However, females also coughed significantly (p = 0.005) more frequently than males: 33 (SD 39) per 30-minute observation and 21 (21) respectively (Fig. S4).

BMI was positively associated with shedding in fine and coarse aerosols in unadjusted models (p < 0.10). BMI was retained in the best fitting adjusted models for both fine and coarse aerosols where it was significantly associated with fine aerosol shedding (p < 0.05). However, BMI was not associated with shedding detected in NP swabs (p > 0.10). Standard categories of BMI were not as good a fit as the continuous BMI and were not significantly associated with shedding (Table S3), although a positive trend is evident for overweight and obese individuals in the adjusted model.

Self-reported vaccination for the current season was associated with a trend (p < 0.10) toward higher viral shedding in fine aerosol samples; vaccination with both the current and previous year’s seasonal vaccines, however, was significantly associated with greater fine aerosol shedding in unadjusted and adjusted models (p < 0.01); In adjusted models we observed 6.3 (95% CI 1.9 to 21.5) times more aerosol shedding among cases with vaccination in the current and previous season compared with having no vaccination in those two seasons. Vaccination was not associated with coarse aerosol or NP shedding (p > 0.10). The association of vaccination and shedding was significant for influenza A (p = 0.03) and not for influenza B (p = 0.83) infections (Table S4).

Viral load in NP swabs was not a significant predictor of aerosol shedding (p = 0.16 for fine and p = 0.48 for coarse aerosols). Temperature measured at the time of sampling, asthma history, smoking, and influenza type were not significantly associated with the extent of measured shedding. While self-reported symptoms were not associated with aerosol shedding, they were significantly associated with shedding measured by the NP swab; only upper respiratory symptoms remained significant when adjusted for other symptoms and age. Increasing age was associated with a significant decrease in shedding in the NP swab; however, age was not associated with aerosol shedding.

## Discussion

We recovered infectious influenza virus from 52 samples of fine aerosols collected from exhaled breath and spontaneous coughs produced by 142 cases of symptomatic influenza infection during 218 clinic visits. Finding infectious virus in 39% of fine aerosol samples collected during 30 minutes of normal tidal breathing in a large community based study of confirmed influenza infection clearly establishes that a significant fraction of influenza cases routinely shed infectious virus, not merely detectable RNA, into aerosol particles small enough to remain suspended in air and present a risk for airborne transmission. Because these data were collected without volunteers having to breathe through a mouthpiece or perform forced coughs, they allow us to provide estimates of average shedding rates, variability and time course of and risk factors for shedding that can be used to provide well-grounded parameter estimates in future models of the risk of airborne influenza transmission from people with symptomatic illness.

The first published estimates of the numbers of influenza virus variants transmitted from donor to recipient host indicated that the bottleneck for transmission between humans is fairly wide and highly variable (mean 192 with 95% confidence 66 – 392).(14, 15) Our observation that cases shed considerable quantities of virus into aerosols, geometric mean >10^4^ RNA copies / 30 minutes and up to 10^3^ infectious virus particles/ 30 minutes, suggests that large numbers of variants could be transmitted via aerosols, especially via the short-range mode.(16) However, longer range aerosol transmission, as might be observed in less crowded environments than in the initial report from Hong Kong, would be expected to usually result in lower exposures and transmission of fewer variants, consistent with the narrower bottleneck described in ferret models.(17)(18)

Sobel et al.(14) suggested that the width of the bottleneck increased with severity of illness, as indicated by a borderline significant positive association between temperature and number of variants transmitted. We did not see a significant association between measured temperature and shedding by any route. In contrast, symptoms were not a significant predictor of bottleneck size, and in our data, symptoms were not significant predictors for shedding into aerosols. Symptoms were, however, significant predictors for nasal shedding as measured in NP swabs. Thus, if aerosols were the more important route of transmission, our observations would be consistent with the currently available bottleneck analysis.

We observed that influenza cases rarely sneezed, despite having just undergone two NP swab collections (a procedure that generally makes one feel an urge to sneeze). Sneezing was not observed in the absence of cough and was not associated with greater aerosol shedding than we observed with cough alone (Fig. S3). Thus, sneezing does not appear to make an important contribution to influenza virus shedding in aerosols. Sneezing might make a contribution to surface contamination. Because sneezes generate considerable amounts of large droplet spray composed of many ballistic droplets not collected by our sampler, we cannot assess that possibility with our data.

Cough was prevalent and was a strong predictor of virus shedding into both coarse and fine aerosols. However, cough was not necessary for infectious aerosol generation in the ≤ 5 μm (fine) aerosol fraction; we detected culturable virus in fine aerosols during 48% of sampling sessions when no coughs were observed. This suggests that exhaled droplets, generated by mechanisms other than cough, are responsible for a portion of the viral load observed in the fine aerosol fraction. Several researchers have recently shown that exhaled aerosol particles are frequently generated from normal healthy lungs by small airway closure and reopening.(19–21) It has been hypothesized that during respiratory infections, airway closure and reopening frequency would be increased due to inflammation with a commensurate increase in aerosol generation and contagiousness.(22)

Cough is thought to produce aerosols from large airways by shear forces that produce relatively coarse aerosol droplets.(23) Our finding that only 13% of cases not observed to have coughed during sample collection produced detectable viral RNA in their coarse aerosols is consistent with that hypothesis. The remaining aerosols may have resulted from speaking; each subject was required to recite the alphabet three times. One might expect that viral replication in the large airways combined with cough generated coarse aerosol droplets would produce the majority of viral aerosols. However, we observed a weak correlation of coarse aerosol RNA copy number with cough frequency and a much stronger association of fine aerosol copy number with cough frequency even though cough would be expected to be the primary source of coarse aerosols. These observations suggest that cough is, at least in part, an epiphenomenon, more of a response to irritation associated with high viral loads in distal airways than a direct source of infectious aerosols.

A striking finding was the association of gender with shedding into fine aerosols. This relationship appears to have resulted from a 3-fold greater impact of coughing on shedding in males. We observed these gender and gender by cough interaction effects only for the fine aerosol fraction. Absence of a gender effect in the coarse aerosol fraction suggests that this is not an effect of cough on aerosol generation by shear forces in the upper airway. We did not measure lung volumes and therefore cannot control for a lung size effect. An equally plausible explanation may be that women tend to have more sensitive cough reflexes.(24) Thus, women may have tended to cough in response to lower viral loads and coughed more frequently at a given viral load, which could have produced the observed steeper slope of viral load regressed on cough frequency in males compared with females. Consistent with this suggestion, we did observe a significantly greater cough frequency in females (p=0.005) and a steeper slope of fine aerosol viral RNA with cough in males (Fig. S4).

BMI was a borderline significant predictor of aerosol shedding in most models, was retained as an important predictor of both coarse and fine aerosols in adjusted models, and reached statistical significance for fine aerosols when adjusted for other factors; it was not a significant predictor of nasal shedding. This observation might be consistent with reports of increased inflammation in models of obesity and influenza and severity of ILI in obese persons. (25–30) Alternatively, increasing BMI is associated with increased frequency of small airways closure and the resulting increased aerosol generation during airway opening as described above may explain the stronger association of BMI with fine than coarse aerosols and lack of association with NP swabs. (31)

Our analysis found a clear separation of factors associated with shedding from the nose and those with shedding into aerosols, especially fine particle aerosols. Upper airway symptoms, as would be expected, were strongly associated with shedding detected in NP swabs, and greatly reduced the size and significance of lower respiratory and systemic symptoms in the fully adjusted model. Age was negatively associated with nasal shedding but not a predictor of aerosol shedding. More surprisingly, no symptoms, including lower respiratory and systemic systems, were strongly associated with shedding into aerosols, in this population with relatively mild lower respiratory symptoms (Fig. 1). Furthermore, nasal shedding was not a significant predictor of aerosol shedding and none of the strong predictors of aerosol shedding were associated with nasal shedding. Thus, we can conclude that the head airways made a negligible contribution to viral aerosol generation and that viral aerosols represent infection in the lung. Moreover, upper and lower airway infection appear to behave as though infection is compartmentalized and independent. In this context, it is notable that Varble et al.(18) observed that intra-host viral variants differ in the nasopharynx and lung of ferrets.

We did not observe a significant decline of viral load detected in NP swabs. If day 1 after onset of symptoms (used as baseline for these analyses) in our cases was equivalent to a mixture of day 1 and day 2 after experimental influenza virus inoculation in the report by Hayden et al.,(32) then our lack of finding a clear drop in nasal shedding over the next two days is reasonably consistent with the pattern reported for experimental infection. There is no available data for comparison of aerosol shedding from published experimental infections. That we saw a much clearer pattern of rapid decline in aerosol shedding again suggests a separation of infection into upper and lower airway compartments in humans.

The association of current and prior year vaccination with increased shedding of influenza A might lead one to speculate that certain types of prior immunity promote lung inflammation, airway closure and aerosol generation. This first observation of the phenomenon needs confirmation. If confirmed, this observation together with recent literature suggesting reduced protection with annual vaccination would have implications for influenza vaccination recommendations and policies.

## Materials and Methods

### Study Population and Sample Collection Procedures

We recruited volunteers with acute respiratory illness on the University of Maryland-College Park campus (UMD) and surrounding community from December 2012 through March 2013.

The UMD Institutional Review Board approved the study, and we obtained a signed consent (or assent and parental verbal assent) from volunteers who reported fever with a cough or sore throat. (Supplemental Fig. S5.)

During the initial visit, we administered a brief screening questionnaire, measured oral temperature, height, weight, and collected two nasopharyngeal (NP) swabs [Copan, Murrieta, CA] for each volunteer screened. One swab was used to perform QuickVue A/B rapid tests for influenza (except when results of a rapid test performed by medical provider were available).

The second NP swab was used for viral culture and PCR for those meeting enrollment criteria and for PCR in a random sample of 24 of those not enrolled.

Participants were asked about sex, age, antipyretic use, vaccination status, use of steroid medications, medical and smoking history, to rate current symptoms on a 4–level scale [none=0, mild=1, moderate=2, severe=3], and to rate the worst symptoms during the illness thus far. We defined symptoms as upper respiratory (runny nose, stuffy nose, sneezing, sore throat, and earache), lower respiratory (chest tightness, shortness of breath, and cough), and systemic (malaise, headache, muscle/joint ache, fever/sweats/chills, and swollen lymph nodes).

Volunteers were enrolled in exhaled breath collection if they met the following criteria: (1) positive QuickVue rapid test, or oral temperature >37.8°C plus cough or sore throat, and (2) presented within the first 3 days of symptom onset. Exhaled breath samples were collected using the Gesundheit-II (G-II) human source bioaerosol sampler as previously described.(12, 33) We collected exhaled breath for 30 min while the participant was seated with their face inside of the large open end of a cone shaped inlet for the G-II. The inlet cone draws in 130 L of air per min and allowed participants to breathe, talk, cough, and sneeze naturally throughout sample collection while maintaining >90% collection efficiency for exhaled and coughed droplets ≤ 100 μm. Subjects were asked to breathe normally and to recite the alphabet once at 5, 15, and 25 min). We collected “coarse” (>5 μm) aerosol droplets by impaction on a Teflon^®^ surface and “fine” droplets (≤5μm and >0.05μm) by condensation growth and impaction on a steel surface constantly rinsed into a buffer containing (phosphate buffered saline with 0.1% bovine serum albumin) liquid reservoir. Audible spontaneous coughs and sneezes during breath collection were counted by direct observation in real-time (n = 59) or by playback of digital recordings (n = 159).

Participants enrolled prior to the third day after symptom onset were asked to come in for up to two consecutive daily follow-up visits (Fig. S5) with repeat questionnaire, NP swab and exhaled breath collections. Final analyses included only visits for enrolled cases occurring on days 1 to 3 post symptom onset with complete data on cough and sneeze, symptoms, PCR results for swab and aerosol samples.

### Laboratory Methods

Detailed methods are described in the supporting information. Briefly, NP swabs were eluted in 1 mL of a phosphate buffered saline with 0.1% bovine serum albumen (PBS/0.1% BSA) or universal transport medium [UTM, Copan, Murrieta, CA], and Teflon^®^ impactors were scrubbed with a nylon swab saturated with PBS/0.1% BSA. The swab was eluted in 1 ml PBS/0.1% BSA. Fine aerosol samples were concentrated to 1 mL using centrifugal ultrafiltration.

RNA was extracted from NP swab, fine and course aerosol samples, and whole-virion standards using an automated Qiagen system and viral RNA was quantified by one-step real-time RT-PCR using Taqman primer probe sets designed by the U.S. Centers for Disease Control and made available through our cooperative agreement. Standard curves were calibrated for virus copy number using plasmids containing a cDNA copy of the RT-qPCR target amplicon. Experimentally determined limits of detection and quantification for each of the RT-qPCR reactions are shown in Table S5.

Virus culture on Madin-Darby canine kidney (MDCK) cells was used to detect infectious virus in NP swab and fine aerosol samples. Coarse aerosol samples were not cultured for infectious virus because impaction on a dry Teflon^®^ surface was expected to reduce infectivity of those samples. Infectious influenza virus was quantified using an immunofluorescence assay for influenza nucleoprotein, and positive cells were counted as FFU by fluorescence microscopy. Details of laboratory methods can be found in the Online Supplement.

### Statistical Analysis

We entered and cleaned data using locally hosted REDCap data capture tools(34) and performed data management and analyses in R (version 3.2.3 R Development Core Team, Vienna, Austria) and SAS (version 9.4, Cary, NC, USA), and produced graphics with Prism Software ((PRISM software version 7.0; GraphPad). We used the delta method to estimate confidence limits for sensitivity and specificity. We used Spearman correlation, generalized linear models (SAS Proc GENMOD), and Tobit regression(35) with nested random effects of sample within subject in (SAS Proc NLMIXED) to analyze infectious virus counts, RNA copy numbers and compute geometric mean virus concentrations. Tobit regression accounted for uncertainty and censoring of the observations by the limit of quantification. We included all independent variables with unadjusted p < 0.10 in initial adjusted models and selected final models using the Akaike information criterion while retaining adjustment for age and sex. Regression model results are presented as the ratio of shedding at the 75^th^ percentile to shedding at the 25^th^ percentile of the distribution of the independent variable so that clinical and epidemiological meaning of the relationship can be more easily interpreted.

## Acknowledgements

This work was funded by US CDC, Cooperative Agreement 1U01P000497 and by NIH grant 5RC1AI086900. The findings and conclusions in this report are those of the authors and do not necessarily represent the official position of the funding agency.

We thank Chengsheng Jiang, Jing Zhang, and Shuo Chen for programming and statistical consultations and the other faculty, staff, and undergraduate research assistants, listed in the supplementary information, who contributed many hours to support this study.

## References

1. Brankston G, Gitterman L, Hirji Z, Lemieux C, Gardam M (2007) Transmission of influenza A in human beings. Lancet Infect Dis 7(4):257–265.

2. Lemieux C, Brankston G, Gitterman L, Hirji Z, Gardam M (2007) Questioning aerosol transmission of influenza. Emerg Infect Dis 13:173–175.

3. Tellier R (2006) Review of Aerosol Transmission of Influenza A Virus. Emerg Infect Dis 12(11):1657–1662.

4. Tellier R (2009) Aerosol transmission of influenza A virus: a review of new studies. J R Soc Interface R Soc 6 Suppl 6:S783–790.

5. Weinstein RA, Bridges CB, Kuehnert MJ, Hall CB (2003) Transmission of Influenza: Implications for Control in Health Care Settings. Clin Infect Dis 37(8): 1094–1101.

6. Killingley B, Nguyen-Van-Tam J (2013) Routes of influenza transmission. Influenza Other Respir Viruses 7:42–51.

7. Aiello AE, et al. (2010) Research findings from nonpharmaceutical intervention studies for pandemic influenza and current gaps in the research. Am J Infect Control 38(4):251–258.

8. Atkinson MP, Wein LM (2008) Quantifying the Routes of Transmission for Pandemic Influenza. Bull Math Biol 70(3):820–867.

9. Nicas M, Jones RM (2009) Relative contributions of four exposure pathways to influenza infection risk. Risk Anal Off Publ Soc Risk Anal 29(9):1292–1303.

10. Spicknall IH, et al. (2010) Informing Optimal Environmental Influenza Interventions: How the Host, Agent, and Environment Alter Dominant Routes of Transmission. PLOS Comput Biol 6(10):e1000969.

11. Lindsley WG, et al. (2016) Viable influenza A virus in airborne particles expelled during coughs versus exhalations. Influenza Other Respir Viruses 10(5):404–413.

12. Milton DK, Fabian MP, Cowling BJ, Grantham ML, McDevitt JJ (2013) Influenza virus aerosols in human exhaled breath: particle size, culturability, and effect of surgical masks. PLoSPathog 9(3):e1003205.

13. Lindsley WG, et al. (2010) Measurements of airborne influenza virus in aerosol particles from human coughs. PloS One 5(11):e15100.

14. Sobel Leonard A, Weissman D, Greenbaum B, Ghedin E, Koelle K (2017) Transmission Bottleneck Size Estimation from Pathogen Deep-Sequencing Data, with an Application to Human Influenza A Virus. J Virol:JVI.00171-17.

15. Poon LLM, et al. (2016) Quantifying influenza virus diversity and transmission in humans. Nat Genet 48(2):195–200.

16. Liu L, Li Y, Nielsen PV, Wei J, Jensen RL (2016) Short-range airborne transmission of expiratory droplets between two people. Indoor Air:n/a−n/a.

17. Frise R, et al. (2016) Contact transmission of influenza virus between ferrets imposes a looser bottleneck than respiratory droplet transmission allowing propagation of antiviral resistance. Sci Rep 6:29793.

18. Varble A, et al. (2014) Influenza A virus transmission bottlenecks are defined by infection route and recipient host. Cell Host Microbe 16(5):691–700.

19. Almstrand A-C, et al. (2010) Effect of airway opening on production of exhaled particles. J Appl Physiol Bethesda Md 1985 108(3):584–588.

20. Johnson GR, Morawska L (2009) The mechanism of breath aerosol formation. J Aerosol Med Pulm Drug Deliv 22(3):229–237.

21. Fabian P, Brain J, Houseman EA, Gern J, Milton DK (2011) Origin of exhaled breath particles from healthy and human rhinovirus-infected subjects. J Aerosol Med Pulm Drug Deliv 24(3):137–147.

22. Edwards DA, et al. (2004) Inhaling to mitigate exhaled bioaerosols. Proc Natl Acad Sci U S A 101(50):17383–17388.

23. Chao CYH, et al. (2009) Characterization of expiration air jets and droplet size distributions immediately at the mouth opening. J Aerosol Sci 40(2):122–133.

24. Kastelik JA, et al. (2002) Sex-related Differences in Cough Reflex Sensitivity in Patients with Chronic Cough. Am J Respir Crit Care Med 166(7):961–964.

25. Cocoros NM, Lash TL, DeMaria A, Klompas M (2014) Obesity as a risk factor for severe influenza-like illness. Influenza Other Respir Viruses 8(1):25–32.

26. Zhou Y, et al. (2015) Adiposity and Influenza-Associated Respiratory Mortality: A Cohort Study. Clin Infect Dis 60(10):E49–E57.

27. Kwong JC, Campitelli MA, Rosella LC (2011) Obesity and Respiratory Hospitalizations During Influenza Seasons in Ontario, Canada: A Cohort Study. Clin Infect Dis 53(5):413–421.

28. Braun ES, et al. (2015) Obesity not associated with severity among hospitalized adults with seasonal influenza virus infection. Infection 43(5):569–575.

29. Louie JK, et al. (2009) Factors Associated With Death or Hospitalization Due to Pandemic 2009 Influenza A(H1N1) Infection in California. JAMA 302(17):1896–1902.

30. Park H-L, et al. (2014) Obesity-induced chronic inflammation is associated with the reduced efficacy of influenza vaccine. Hum Vaccines Immunother 10(5): 1181–1186.

31. Salome CM, King GG, Berend N (2010) Physiology of obesity and effects on lung function. J Appl Physiol 108(1):206–211.

32. Hayden FG, et al. (1998) Local and systemic cytokine responses during experimental human influenza A virus infection. Relation to symptom formation and host defense. J Clin Invest 101(3):643–649.

33. McDevitt JJ, et al. (2013) Development and Performance Evaluation of an Exhaled-Breath Bioaerosol Collector for Influenza Virus. Aerosol Sci Technol J Am Assoc Aerosol Res 47(4):444–451.

34. Harris PA, et al. (2009) Research electronic data capture (REDCap)—A metadata-driven methodology and workflow process for providing translational research informatics support. JBiomedInform 42(2):377–381.

35. Twisk J, Rijmen F (2009) Longitudinal tobit regression: A new approach to analyze outcome variables with floor or ceiling effects. J Clin Epidemiol 62(9):953–958.

